# ABA-responsive transcription factor CbNAC63 is involved in modulating biosynthesis of triterpenoid saponins in *Conyza blinii*

**DOI:** 10.1101/2025.11.27.690938

**Authors:** Yizhen Yuan, Min Zhou, Ming Yang, Maojia Wang, Junyi Zhan, Tao Wang, Hui Chen, Tianrun Zheng

## Abstract

*Conyza blinii*, a traditional Chinese medicinal herb, accumulates triterpenoid saponins. NAC transcription factors (TFs) are pivotal for plant stress responses and hormone-mediated secondary metabolism, while abscisic acid (ABA) regulates terpenoid biosynthesis in medicinal plants. However, how NAC TFs mediate ABA-regulated triterpenoid saponin synthesis in *C. blinii* remains unclear. In order to address this gap, the focus was directed towards CbNAC63—a NAC TF identified from the nocturnal low-temperature transcriptome of *C. blinii*. Phylogenetic and motif analyses showed it belongs to the stress-responsive NAC subclade with conserved domains. Y2H assays revealed its self-interaction, enhancing transcriptional activity; subcellular localization confirmed nuclear localization. RT-qPCR results indicated CbNAC63 exhibits tissue-specific expression under nocturnal low-temperature stress, accompanied by increased triterpenoid saponin accumulation. ABA signaling primarily regulates this process: exogenous ABA enhanced CbNAC63 expression and saponin content, while fluridone suppressed both. Overexpression (OE) and virus-induced gene silencing (VIGS) demonstrated CbNAC63 positively regulates saponin biosynthesis. Y1H and dual-luciferase reporter assays confirmed CbNAC63 directly binds to the promoters of *Cb*β*AS* and *CbHMGR* and activates their transcription. In summary, the study shows CbNAC63 integrates ABA and low-temperature signals, activating MVA pathway genes to regulate saponin synthesis. This unveils a novel “ABA-CbNAC63-MVA” module, enriches the classic regulatory network, and provides a target for improving medicinal plant quality.

**Highlights:** Identifies CbNAC63 as a novel regulator that integrates ABA and cold signals to boost saponin production in medicinal plants.

## Introduction

As sedentary organisms, plants cannot avoid or migrate in the same way as animals when they are exposed to abiotic stresses in the external environment(Zhang et al., 2021). Plants are vulnerable to environmental stresses such as temperature, heavy metals, salt stress, etc(Gong et al., 2020). To resist environmental stress, plants can only improve their adaptability through their own physiological processes related to hormones signaling, RNA transcription regulation, protein translation and modification(Waadt et al., 2022, Mizoi et al., 2011). Environmental temperature is an influential factor governing plant growth and development, plants are normally active growth only at optimum temperature(Sakamoto and Kimura, 2018). Although the adverse environment is not beneficial to crop reproductive growth and increased food production, it is a beneficial factor in improving the quality of medicinal plants.

“Environmental factors - phytohormones - transcription factors - secondary metabolites” regulatory networks have been reported in a range of medicinal plants, such as *Artemisia annua* L., *Salvia miltiorrhiza* Bge., *Taxus* spp., *Catharanthus roseus* L., involving active ingredients such as terpenoids, alkaloids, saponins and phenolic acids, etc(Zheng et al., 2023). Jasmonate-induced AaERF1, AaERF1 and AaMYC2 can be positively factors regulate artemisinin biosynthesizing within *Artemisia annua*(Zong Xia et al., 2011, Shen et al., 2016). TcWRKY33 positively regulates taxoid biosynthesis by transmitting salicylic acid (SA) signaling(Chen et al., 2021) . Downstream of the SA signaling pathway, AaTGA6 regulates artemisinin synthesis by binding to the *AaERF1* promoter via the AaNPR1/AaTGA3 module(Zongyou et al., 2019). Overexpression of the ABA receptor orthologue, AaPYL9 in *A. annua* increases not only drought tolerance, but also improves artemisinin biosynthesis(Zhang et al., 2013). SmbZIP1, a basic leucine zipper transcription factor was separated from ABA-induced transcriptome, which positively promotes the phenolic acid biosynthesis(Deng et al., 2020). In *Taxus,* the expression of *TcMYB29a* can be upregulated by ABA, then binds with the promoters of taxol-biosynthesis-related genes and increasing the content of taxol(Cao et al., 2022b).

This shows that tapping the key regulators of secondary metabolites in medicinal plants is one of the cores of genetic engineering methods to enhance the quality of medicinal plants. NAC transcription factors (TFs) family has a variety of important regulatory roles in plants which are related to plant growth, development, fruit ripening, and stress response(Moyano et al., 2018, Liu et al., 2023a, Diao et al., 2020). Admittedly, these regulatory processes are frequently associated with plant hormones(Kou et al., 2021). NAC proteins could form homodimers or interact with other TFs to produce heterodimers(Olsen et al., 2005). The ABA-induced NAC transcription factor MdNAC1 promotes anthocyanin synthesis in red-fleshed apples by interacting with MdbZIP23 to upregulate the expression of MdMYB10(Liu et al., 2023b). The transgenic overexpressing *AaNAC1* in *A. annua* exhibited increased tolerance to drought and resistance to *Botrytis cinerea*(Zongyou et al., 2016). SmNAC1, a UV-B-mediated transcription factor, promoting the synthesis of salvianolic acid by upregulating the expression of *PAL3* and *TAT3* genes(Xiaojian et al., 2020).

*Conyza blinii* (*C. blinii*) is a traditional Chinese medicinal herb distributed in southwest China and its active medicinal ingredient is terpenoid. Triterpenoid saponins are glycosides with remarkable structural and bioactive diversity, becoming increasingly significant in the treatment of cancer due to their efficacy and safety(Yao et al., 2020, Haralampidis et al., 2002). The medicinal ingredient triterpenoidal saponins of *C. blinii* has been proven to have an anti-cancer effect by inhibiting the autophagy of Hela cells(Liu et al., 2017). β-amyrin synthase (βAS) converts 2,3-oxidosqualene to β-amyrin(Hoshino et al., 2015), which is the basic structure of oleanane-type triterpenoid. In our previous studies, we reported that CbWRKY24 is induced by JA to mediate the accumulation of saponins(Sun et al., 2018). And another CbNAC63 transcription factor, whith potential positive relevance to saponin synthesis has been identified in our previous nocturnal low-temperature transcriptome(Yang et al., 2023).

In this study, we screened that CbNAC63 could be a positive regulator of triterpenoid saponins synthesis in *C. blinii*. Specifically, CbNAC63 enhances the production of triterpenoid saponins by interacting with the promoter of *Cb*β*AS*, especially in the presence of ABA. The findings of this study reflect the NAC and MVA pathway in a new molecular mechanism promoting triterpenoid saponins synthesis in the ABA signaling pathway.

## Materials and methods

### Plant materials

Two-month-old seedlings were selected for experiment. The relative humidity of the laboratory was 50-70%. All of the plants were managed under a long-day photoperiod (16 h: 8 h, light: dark).

### Extraction of saponins and content analysis

The vanillin (5%, mass fraction)-glacial acetic acid colorimetry was employed to quantify saponin content, following the protocol established in our previous study.

### Phylogenetic tree and motif analysis

The phylogenetic tree which based on gene-terpenoid co-expression networks refer to our previous research(Yang et al., 2023). Motif analysis: The output .xmL file was downloaded to the MEME website: (https://meme-suite.org/meme/), and TBtools software was used for motif sequence and phylogenetic tree visual analysis.

### Plant total RNA extraction and RT-qPCR analysis

Total RNA required for this experiment was extracted and reverse transcribed into cDNA by the experimental method mentioned above(Sun et al., 2018, Zheng et al., 2020). *CbGAPDH* was selected as the internal reference gene and the genes expression were calculated by the 2^−ΔΔC^ method. The method of RT-qPCR have been mentioned in our studies(Yang et al., 2022).

### Subcellular localization and transcriptional activation

Subcellular localization and transcriptional activation methods refer to our published research for details(Yang et al., 2023).

### Hairy roots

The cultivation of hairy roots was modified according to the method of Cao et al(Cao et al., 2022a). The pCambia1300-*CbNAC63-*GFP recombinant vector was transferred into K599 receptor cells. Colonies were formed on plates containing Kan and Rif (50mg/ml). The colonies were added to YEB liquid medium cultured overnight and allowed to grow until the OD600 increased to 1. *C. blinii* which has been cut off from the root tissue is immersed in the bacterial solution for a few seconds. Infected above-ground tissue will be inserted into the soil and continued to be cultured until new roots. GFP signals in roots was observed by laser-scanning microscope (Olympus).

### Agrobacterium tumefaciens-mediated transient transformation and virus induced gene silencing (VIGS) experiments

The vector used for agrobacterium-mediated transient overexpression is pCambia1300, and the vectors used for VIGS are pTRV_1_ and pTRV_2_. Experimental methods have been mentioned in our published study(Yang et al., 2023).

### Dual-luciferase reporter gene detection

The recombinant vectors pGreen-62SK-*CbNAC63* and pGreen II-0800-LUC (containing the promoters of *Cb*β*AS* and *CbHMGR*) were introduced into Agrobacterium tumefaciens GV3101 (pSoup) competent cells. Positive transformants were identified by colony PCR. Successfully transformed colonies were used to prepare Agrobacterium suspensions, which were mixed 1:1 in pairs and kept in the dark for 2 h. The mixed Agrobacterium suspensions were then infiltrated into *Nicotiana benthamiana* leaves. After 48–72 h, the infiltrated leaves were excised and 0.1 mM Beetle Luciferin Potassium Salt solution was applied to the abaxial side. Following a 10-min dark incubation, luminescence was visualized with an in vivo imaging system. The remaining leaf samples were processed with the Dual-Luciferase® Reporter Assay Kit to measure LUC and REN activities according to the manufacturer’s instructions.

### Yeast one-hybrid and two-hybrid assay

The recombinant plasmids pGADT7-Rec2-*CbNAC63*, pro*Cb*β*AS*-pHIS2, and pro*CbHMGR*-pHIS2 were constructed and transformed into Y187 yeast competent cells. After transformation, the cells were evenly spread on SD-Trp/Leu solid medium and incubated at 30°C for 48–96 h. Single positive colonies were picked and inoculated into 10 mL of SD-Trp/Leu liquid medium, followed by 24 h of shaking at 30°C. The cultures were centrifuged at 5000 rpm for 10 min, the supernatants were discarded, and the pellets were resuspended in 0.9% NaCl, centrifuged again under the same conditions, and finally resuspended in 0.9% NaCl to an OD600 of approximately 0.8. Ten-fold serial dilutions were prepared, and 5 µL of each dilution was spotted onto SD-Trp/Leu plates as well as onto SD-Trp/Leu/His plates supplemented with 0, 10, 20, or 30 mM 3-AT. After incubation at 30C for 2 d, the growth of the yeast colonies was assessed.

The CDS of CbNAC63 was cloned into both pGADT7 and pGBKT7 vectors. The resulting recombinant plasmids pGBKT7-CbNAC63 and pGADT7-CbNAC63 were co-transformed into AH109 yeast competent cells. Transformed cells were evenly spread on SD-Trp/Leu solid medium and incubated at 30 °C for 2 days. A single colony was picked and inoculated into 10 mL SD-Trp/Leu liquid medium, then cultured at 30 °C with shaking for ∼24 h. The culture was centrifuged at 5000 r/min for 10 min, the supernatant was discarded, the pellet was resuspended in 0.9% NaCl, centrifuged again at 5000 r/min for 10 min, and finally adjusted with 0.9% NaCl to OD600 = 0.2–0.4. Ten-fold serial dilutions were prepared, and 5 μL of each dilution was spotted onto SD-Trp/Leu plates and onto SD-Trp/Leu/His/Ade plates supplemented with 0, 10, 20, or 30 mM 3-AT. After incubation at 30 °C for 2 days, colony growth was assessed. (pGADT7-T + pGBKT7-53 served as the positive control, and AH109 alone as the negative control.)

## Result

### The effect of nocturnal low temperature on the *C. blinii*

In order to investigate the effect of nocturnal low temperature on the *C. blinii,* we performed an analysis of phenotypic changes and quantification of saponin content. As the duration of treatment increased, leaf wilting and curling progressively worsened, with some leaves exhibiting marked morphological deformities and reduced stem support capacity. It’s evidently that nocturnal low temperatures severely inhibited the normal development of *C. blinii* (Figure 1A). RT-qPCR results revealed that *CbHMGR* transcription levels were significantly increased at 1d, while *CbSQS*, *CbFPPS*, and *Cb*β*AS* expression exhibited a transient increase followed by decline over time. This indicated that nocturnal low temperatures regulate transcription of MVA-pathway genes(Figure 1B). Additionally, saponin content markedly increased across all tissues, with oleanolic acid levels in stems consistently far exceeding those in leaves and roots(Figure 1C). This finding suggests that low temperatures trigger the coordinated activation of the terpenoid biosynthesis pathway and saponin accumulation, with stems serving as the primary storage organ for saponins.

**Fig. 1.**
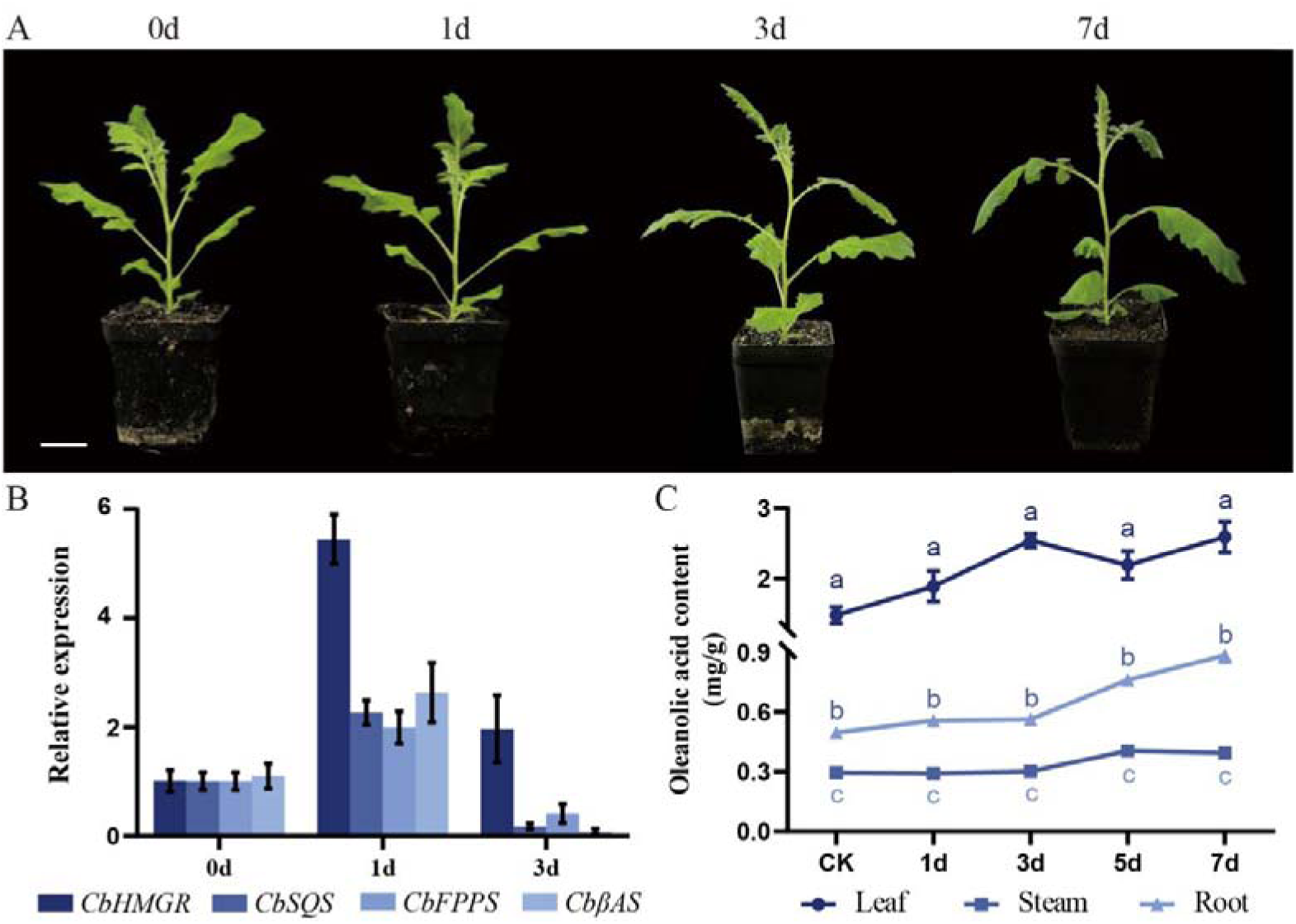
Phenotypic and saponin content changes of *C. blinii* under nocturnal low temperature. (A) The phenotypic changes of *C. blinii*, Bar=2 cm. (B) The changes in the transcript levels of MVA-pathway enzyme genes in *C. blinii.* (C) The tissue-specific variations in saponin content in *C. blinii*. Data values are means ± SD (n=3).

### NAC transcription factor CbNAC63 involved in

Phylogenetic analysis clusters CbNAC63 within the stress-responsive NAC subclade, thus demonstrating that CbNAC63 belongs to the NAC transcription factor family and shares evolutionary relationships with NAC homologs from other species. Conserved motif analysis identified a DNA-binding NAC domain (Motif 1) and a C-terminal regulatory region (Motif 3). It is notable that CbNAC63 shares multiple conserved motifs with NAC proteins from other species, thereby reflecting the evolutionary stability of these motifs. This provides a structural basis for its biological functions and implies that it may mediate core functions analogous to those of other NAC proteins(Figure 2A). In conditions of ambient temperature, the expression of *CbNAC63* in stems exhibited a significantly higher level of expression in comparison to that observed in roots and leaves. During the nocturnal low-temperature, the expression of *CbNAC63* in leaves demonstrated dynamic fluctuations, yet remained significantly elevated in comparison to leaves cultivated under standard temperature conditions. The present indicated that CbNAC63 not only displays tissue-specific expression but also participates in the plant’s response to nocturnal low-temperature stress(Figure 2B). Subsequently, yeast two-hybrid (Y2H) assays further revealed that CbNAC63 forms functional homodimers. (Figure 2C). This study confirmed that CbNAC63 self-interacts to form homodimers, providing crucial molecular evidence for its function as a transcription factor and laying the groundwork for elucidating its role in regulating triterpenoid saponin biosynthesis. Meanwhile, the yeast transcriptional autoactivation assay demonstrated that the transcription factor CbNAC63 possesses transcriptional autoactivation activity and is capable of independently initiating reporter gene transcription in the yeast system(Figure 2D). Subcellular localization confirmed that CbNAC63 exists in the nucleus. This nuclear localisation is consistent with the established function of transcription factors in the regulation of gene expression within the nucleus(Figure 2E).

**Fig. 2.**
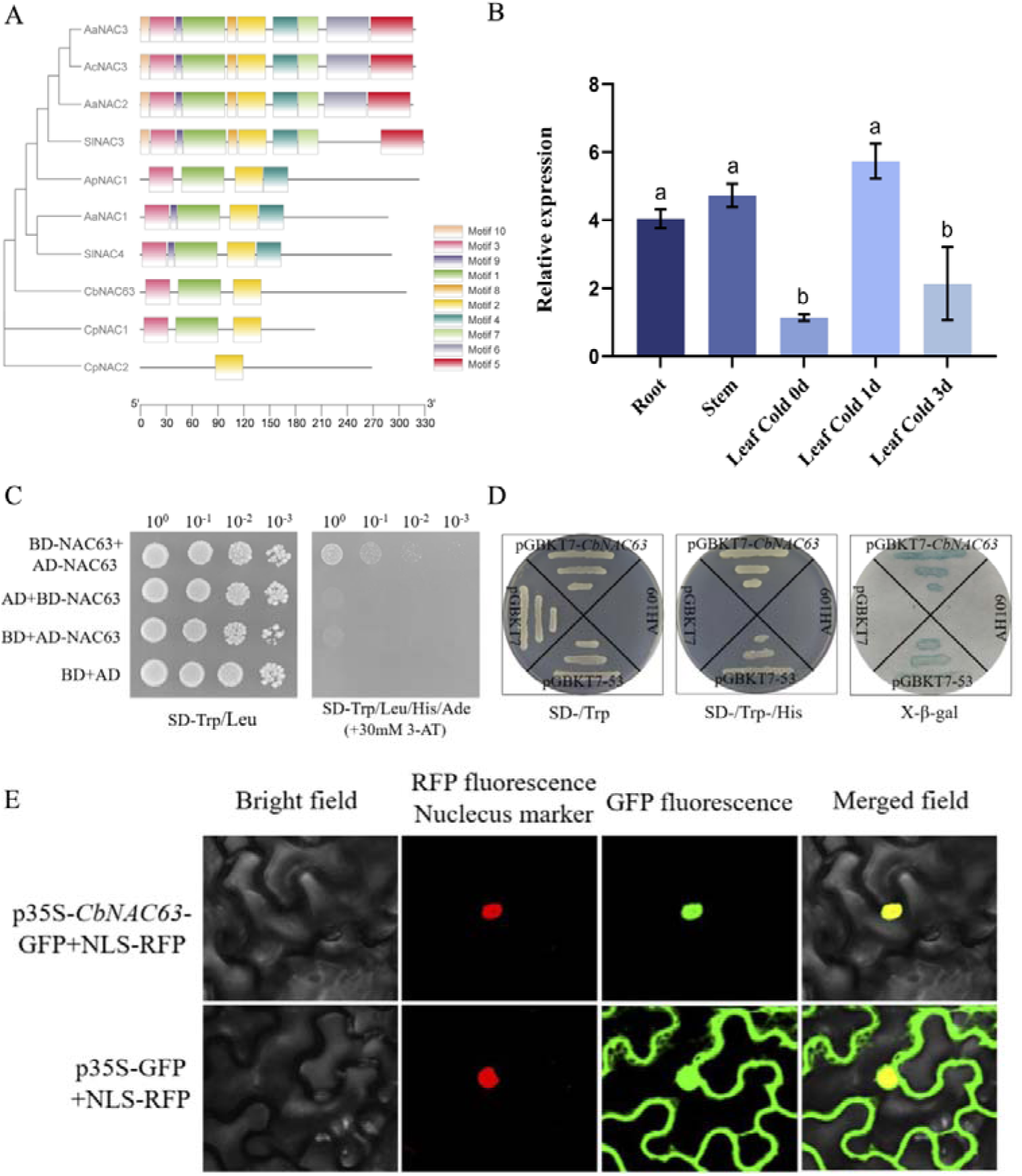
The motif and expression-pattern analyses of the *CbNAC63* transcription factor. (A) The Phylogenetic and motif analyses of CbNAC63. Rectangles rendered in distinct colors denote different yet evolutionarily conserved motifs. (B) RT-qPCR revealed both tissue-specific expression and nocturnal-low-temperature responsiveness of CbNAC63. (C) Yeast two hybrid (Y2H) assay confirms CbNA C63 self-interaction. (D) CbNAC63 Transcriptional autoactivation assay of CbNAC63 in yeast system. (E) Subcellular localization of CbNAC63 by p35S-CbNAC63-GFP. Green: GFP; Red: nuclear localization signal (NLS). Bar=50 µm. Data values are means ± SD (n=3).

### ABA plays a dominant role to affect the gene expression

According to pre-study (life and RNA-seq No.), abscisic acid (ABA) is a phytohormone signal of significance, which can participate in the MVA/MEP pathway to regulate the synthesis of terpenoids, thereby helping plants adapt to environmental changes caused by low temperature stress. To further confirm the role of ABA in affecting gene expression, we examined the expression of ABA-signaling pathway genes by qRT-PCR. It was found that under exogenous treatments, the relative expression levels of ABA signal transduction genes exhibited differential responses. *CbPP2C* and *CbABF* were significantly upregulated, indicating activation of the ABA signaling pathway (Figure 3A-B). For key MVA-pathway enzymes and *CbNAC63*, their transcript abundances in leaves and roots were regulated by exogenous ABA or fluridone (FDT) in a time-dependent and tissue-specific manner, with *CbNAC63* expression peaking at 48 hours as indicated by heatmap profiles (Figure 3C). Furthermore, exogenous ABA and fluridone (FDT) treatments influenced saponin content in *C. blinii* leaves. The effects of ABA treatment on saponin levels were found to be both duration-dependent and application-method-dependent, whereas saponin accumulation was reduced in FDT-treated groups (Figure 3D-E). This suggests that that ABA signaling plays a crucial role in regulating saponin biosynthesis and accumulation.

**Fig. 3.**
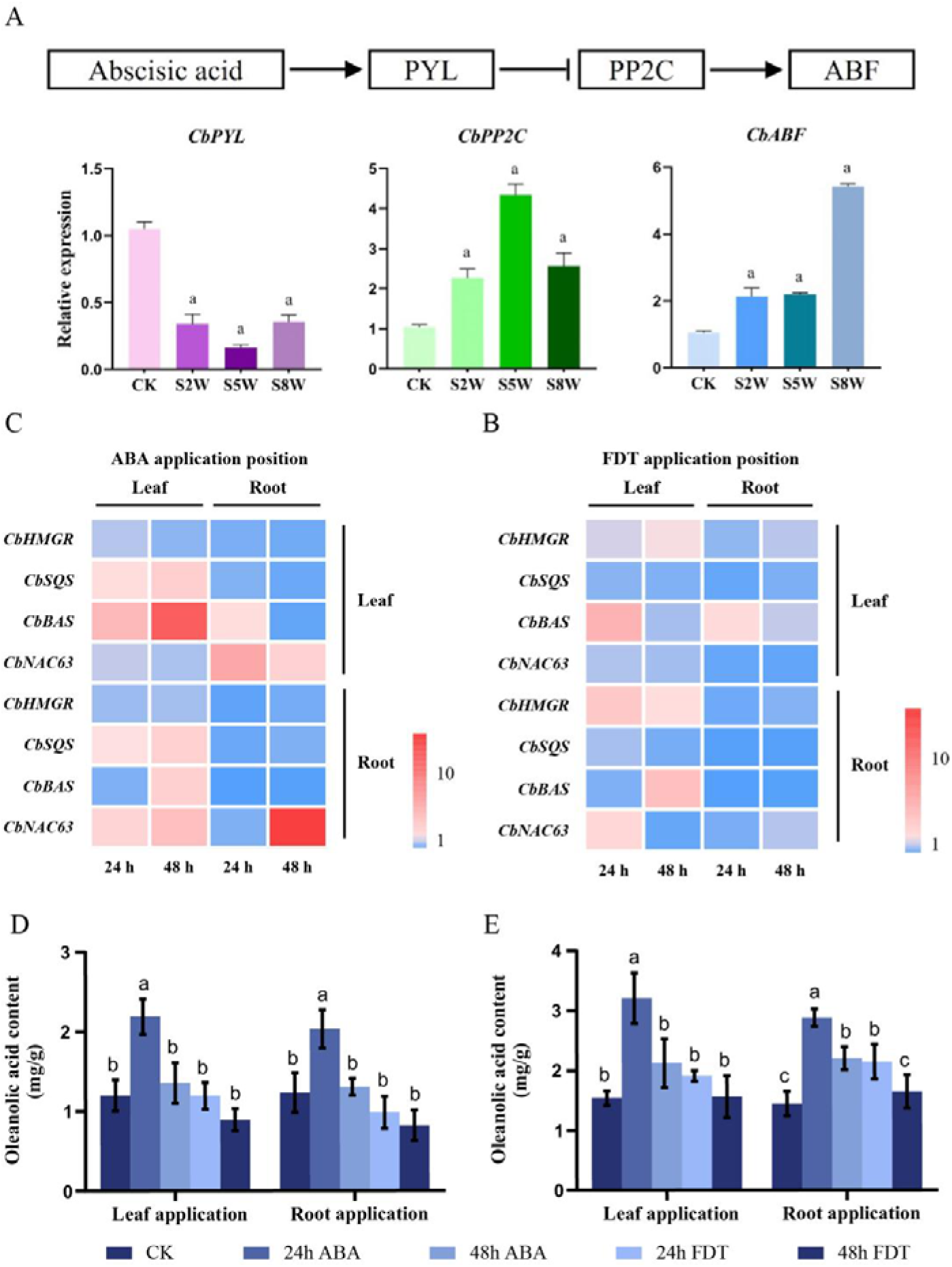
ABA signal modulates both CbNAC63 expression and saponin accumulation in *C. blinii*. (A) Relative expression levels of ABA signal-transduction genes (left to right: *CbPYL*, *CbPP2C*, and *CbABF*). (B-C) Heat-map showing the relative transcript abundance of key MVA-pathway enzymes and *CbNAC63* in leaves and roots after 24h and 48h of exogenous ABA or Fluridone (FDT, ABA biosynthesis inhibitor) treatments; color intensity represents gene-expression values. (D) Effect of exogenous ABA and fluridone (FDT) on saponin content in *C. blinii* leaves after 24h and 48h treatments with root-drench and leave-spray. (E) Effect of exogenous ABA and fluridone (FDT) on saponin content in *C. blinii* leaves after 24h and 48h treatments with leave-spray. Data values are means ± SD (n=3).

### CbNAC63 functions as a key regulator of saponin content

The recombinant plasmid pCambia1304-*CbNAC63*-GFP was introduced into *Agrobacterium rhizogenes* strain K599 by electroporation. Freshly cut root–stem junctions of *C. blinii* seedlings were immersed in the K599 suspension (Figure 4A). Following a culture period of 4–6 weeks, observation under a laser-scanning confocal microscope revealed that the green fluorescence colocalized with root tissues (Figure 4B-D), confirming the successful expression and localized of CbNAC63 in hairy roots. Subsequently, the GFP-positive hairy roots were then harvested for downstream analyses. In order to clarify the regulatory role of CbNAC63 in saponin biosynthesis, overexpression and virus-induced gene silencing experiments were conducted. The results of the experiment demonstrated that the expression of CbNAC63 overexpression resulted in a marked elevation of its transcript levels in hairy roots. In contrast, VIGS-mediated silencing reduced *CbNAC63* transcription by about 70% (Figure 4E-G). The content of saponins also showed the same change. The hairy root lines with *CbNAC63* overexpression accumulated 2.2-fold more saponins in comparison to the control, while the saponins levels in the silenced lines were reduced by about 40% compared with the control (Figure 4H-J). The results obtained demonstrate unequivocally that the transcription factor CbNAC63 exerts a positive regulatory influence on saponin biosynthesis and accumulation in *C. blinii*.

**Fig. 4.**
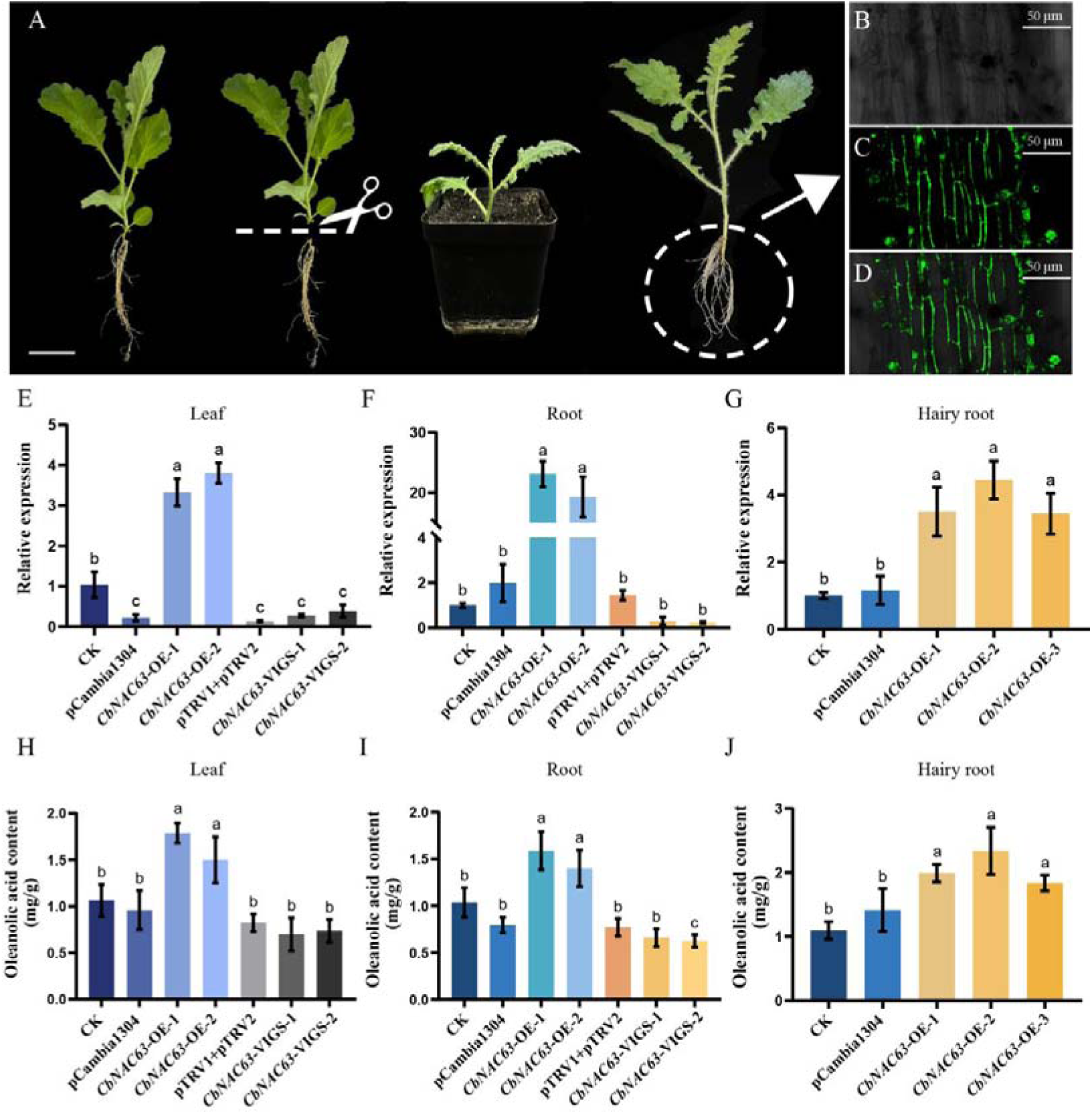
Overexpression and silencing experiments demonstrate that *CbNAC63* modulates saponin accumulation. (A) Non-sterile hairy-root transformation protocol. Recombinant plasmid pCambia1304-*CbNAC63*-GFP was introduced into *Agrobacterium rhizogenes* strain K599 by electroporation. Freshly cut root–stem junctions of *C. blinii* seedlings were immersed in the K599 suspension. After 4–6 weeks of culture, GFP-positive hairy roots emerging at the wound site were harvested for downstream analyses. Bar=2 cm. (B-D) Positive hairy roots were identified under a laser-scanning confocal microscope. (B) Bright-field; (C) dark-field; (D) merged channel. Green: GFP (green fluorescent protein). Bar=50 μm. (E-G) RT-qPCR quantification of *CbNAC63* transcript levels in transiently transformed *C. blinii* tissues: leaves, roots, and hairy roots. (H-J) Changes in saponin content across transiently transformed *C. blinii* tissues - leaves, roots, and hairy roots. Data values are means ± SD (n=3).

### CbNAC63 modulates the transcription of *Cb***β***AS* and *CbHMGR* by directly binding to their promoters

In order to elucidate the specific mechanismof CbNAC63 in regulating triterpenoid saponin biosynthesis, we performed RT-qPCR to quantify the expression of key MVA pathway genes in CbNAC63-overexpressing hairy roots. In comparison with the control group, OE lines demonstrated a substantial upregulation of *Cb*β*AS* and *CbHMG*R. CbNAC63 silencing reduced their expression to 0.3-0.5-fold of control levels (Figure 5A, C). In contrast, other MVA pathway genes (*CbFPPS, CbSQE, CbSQS*) showed no response to overexpression, yet demonstrated modest downregulation in silenced lines though less pronounced than *Cb*β*AS/CbHMGR* (Figure 5B, D, E). These results demonstrate that CbNAC63 selectively enhances the expression of rate-limiting enzymes (*Cb*β*AS* and CbHMGR) while exerting minimal effects on downstream MVA genes.Y1H results indicate that yeast simultaneously transfected with CbNAC63 and either the *Cb*β*AS* or *CbHMGR* promoter grew normally on SD-Trp/Leu/His selective medium supplemented with 20 mM 3-AT, whereas the control lacking CbNAC63 promoter-specific binding exhibited growth restriction. This confirms that CbNAC63 directly binds to both *Cb*β*AS* and *CbHMGR* promoters in vitro. To further validate this binding occurs in vivo, a dual luciferase reporter assay was performed. CbNAC63 strongly induced luciferase (LUC) activity driven by p*roCb*β*AS*. Compared to the control group, the *proCb*β*AS* + CbNAC63 group exhibited significantly higher fluorescence intensity, with the LUC/REN ratio increasing more than 3.5-fold (P < 0.0001; Figure 5I-J). Similarly, CbNAC63 activated *proCbHMGR*. LUC fluorescence intensity in the p*roCbHMGR*+CbNAC63-SK group exceeded that of the control group, with a significantly elevated LUC/REN ratio (P < 0.001; Fig. 5K-L). These in vivo results unequivocally confirm that CbNAC63 specifically transactivates *Cb*β*AS* and *CbHMGR*.

**Fig. 5.**
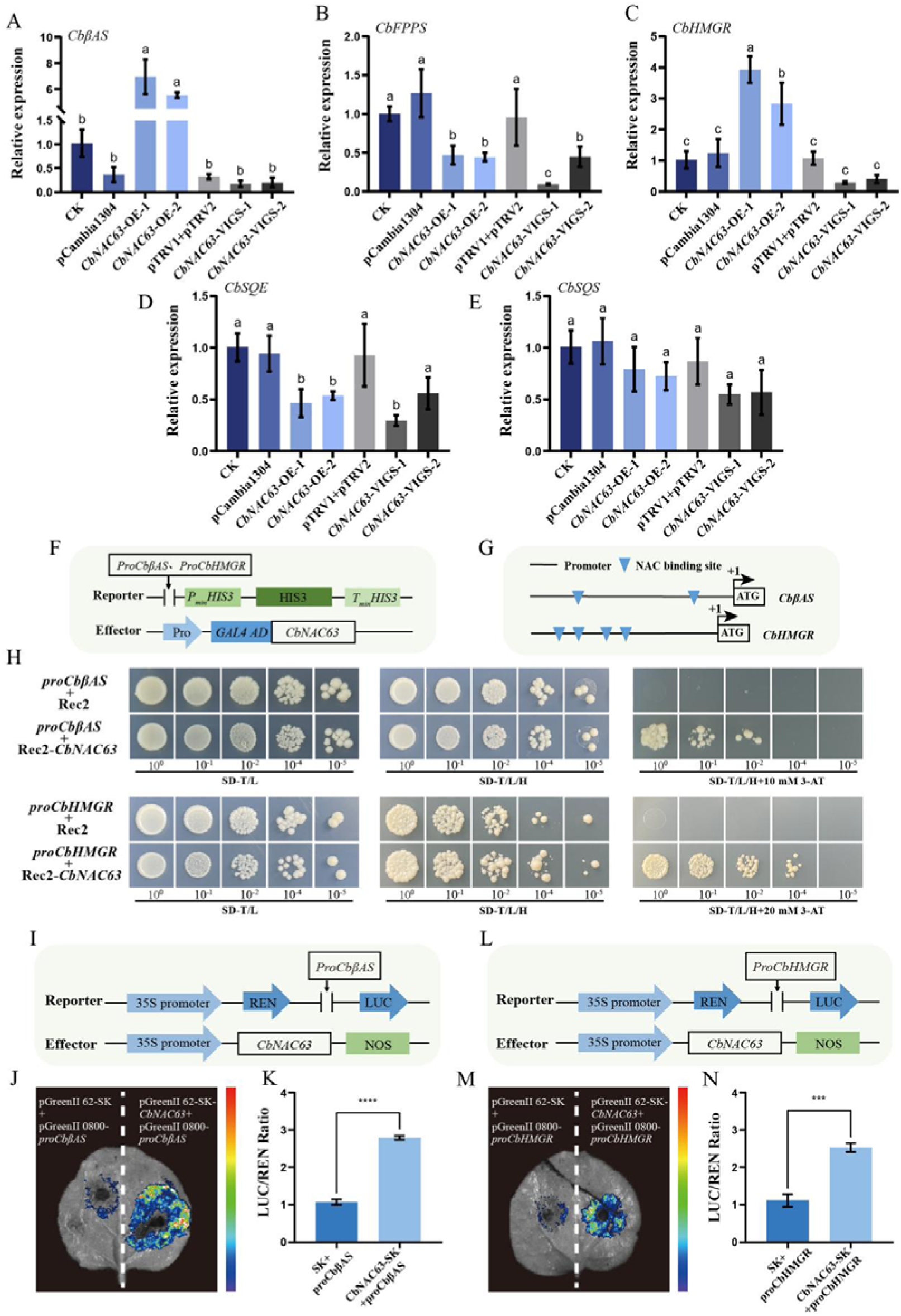
CbNAC63 participates in the transcriptional regulation of the *Cb*β*AS*. (A-E) RT-qPCR quantification of key MVA-pathway enzyme genes in *CbNAC63*-overexpressing hairy roots. (F, I and L) Schematic diagrams of the reporter and effector constructs. (G) Analysis of NAC-binding sites in the promoters of *Cb*β*AS* and *CbHMGR*. (H) Yeast one-hybrid (Y1H) assays confirm the direct binding of CbNAC63 to the *Cb*β*AS* and *CbHMGR* promoters. (J and M) Dual-luciferase reporter assay images showing CbNAC63-mediated activation of *proCb*β*AS* (J) and *proCbHMGR* (M). (K and N) Dual-luciferase reporter assay (LUC/REN ratios) for CbNAC63 with *proCb*β*AS* (K) and *proCbHMGR* (N). ****: P<0.0001, ***: P<0.001. Data values are means ± SD (n=3)

**Fig.6.**
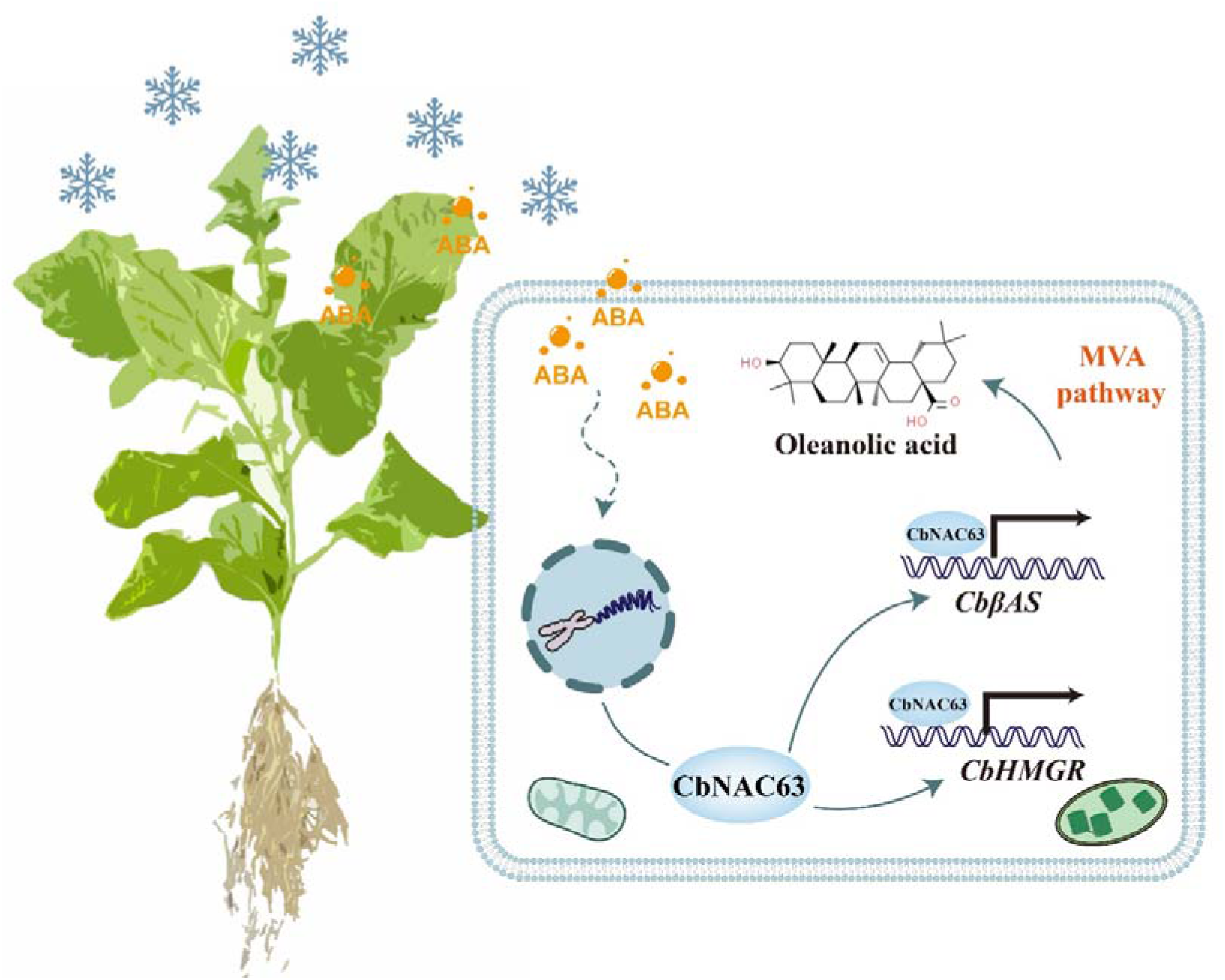
‘ABA signal-*CbNAC63*-*CbHMGR*/*Cb*β*AS*-Ola’ a feedback-approach to escape from nocturnal low-temperature

## Discussion

This study focuses on the regulatory mechanisms governing triterpenoid saponin synthesis in *C. blinii*. The core findings reveal that the ABA-responsive transcription factor CbNAC63 directly activates the expression of key MVA pathway genes *Cb*β*AS* and *CbHMGR*, thereby promoting triterpenoid saponin biosynthesis in *C. blinii* under nocturnal low-temperature stress. This study establishes the ‘ABA-CbNAC63-MVA’ axis as a core mechanism integrating hormonal and environmental signals to enhance secondary metabolite accumulation.

ABA can act synergistically with jasmonic acid (JA) and salicylic acid (SA) to fine-regulate secondary metabolism. In *Artemisia annua*, ABA and JA cooperatively enhance artemisinin biosynthesis through the synergistic activation of AaMYC2 and AaERF1/2(Shen et al., 2016; Zong Xia et al., 2011). Similarly, SA signaling in Taxus chinensis recruits TcWRKY33 to upregulate taxol biosynthetic genes, a process potentiated by ABA(Chen et al., 2021). Our data extend this paradigm to *C. blinii*, where ABA-induced CbNAC63 activation coincides with JA-mediated CbWRKY24 upregulation(Sun et al., 2018). This suggests hormonal crosstalk (ABA/JA/SA) converges on TF networks to amplify saponin production. Future studies should dissect whether CbNAC63 physically interacts with JA/SA-responsive TFs (e.g., CbWRKY24) to form heterodimeric complexes, as observed in apple anthocyanin synthesis(Liu et al., 2023b).

CbNAC63 as a terminal TF: Direct target binding and potential upstream hierarchical regulation. Our data confirm CbNAC63 directly binds *Cb*β*AS* and *CbHMGR* promoters, implying it acts as a terminal TF for these genes. However, promoter analyses of homologous NAC targets reveal ABA-responsive elements (ABREs) that recruit upstream regulators(Cao et al., 2022b). Whether CbNAC63 itself requires activation by ABA-induced TFs remains untested. In *Salvia miltiorrhiza*, SmbZIP1— an ABA-inducible TF—binds NAC gene promoters to regulate phenolic acids(Deng et al., 2020), suggesting hierarchical TF cascades may operate upstream of CbNAC63. Additionally, CbNAC63 forms homodimers, raising the possibility of heterodimerization with other TFs to expand target gene specificity.

Triterpenoid saponins in *C. blinii* are predominantly oleanane-type, with oleanolic acid (Ola) as the core aglycone. Oleanolic acid, as the core skeleton of pentacyclic triterpenoid compounds, exhibits metabolic regulation, organ protection, anti-inflammatory, antioxidant, and antitumor activities(Liu, 1995). Low-temperature stress significantly increased Ola accumulation in stems, aligning with its role in stress adaptation. Ola derivatives exhibit potent anticancer activity by inducing apoptosis and inhibiting autophagy in cancer cells(Liu et al., 2017). Notably, structural modifications enhance bioactivity; for example, blinin saponins from *C. blinii* inhibit HeLa cell autophagy via dual-targeting mTOR and Beclin-1 pathways(Liu et al., 2017). Our study confirms ABA and CbNAC63 elevate total saponins, but whether they alter Ola derivative profiles warrants metabolomic interrogation.

### Conclusions

In conclusion, we observed CbNAC63, a stress-responsive NAC transcription factor in *C. blinii*, as a key regulator of triterpenoid saponin biosynthesis. CbNAC63 integrates ABA and low-temperature signals, directly binding to the promoters of key MVA pathway genes *Cb*β*AS* and *CbHMGR* to activate their transcription, thereby positively regulating triterpenoid saponin synthesis in *C. blinii*. This study first reveals the ‘ABA-CbNAC63-MVA’ regulatory module, enriching the ‘environment-hormone-transcription factor -secondary metabolism’ regulatory network in medicinal plants and providing a novel case study for understanding the functional diversity of NAC family transcription factors.

## Acknowledgments

We extend our gratitude to all colleagues in the laboratory for their valuable discussions and technical assistance. Special thanks are due to the editors and reviewers for their rigorous assessment of the manuscript and for their constructive suggestions for improvement.

## Author Contributions

T.Z. : conceptualization; M.W., and M.Z.: visualization; T.Z., and M.Y.: formal analysis; M.Z., and M.W.:investigation; Y.Y., M.Z., and M.W.: data curation; Y.Y.: writing - original draft; Y.Y., T.Z.: writing - review & editing; T.Z., M.Z., T.W., and H.C.: supervision; T.Z., and H.C.: funding acquisition. All authors have read and agreed to the published version of the manuscript.

## Data Availability Statement

Not applicable.

